# Metabolic labelling of RNA uncovers the contribution of transcription and decay rates on hypoxia-induced changes in RNA levels

**DOI:** 10.1101/694570

**Authors:** Maria Tiana, Clara Galiana, Miguel Ángel Fernández-Moreno, Benilde Jimenez, Luis del Peso

## Abstract

Cells adapt to environmental changes, including fluctuations in oxygen levels, through the induction of specific gene expression programs. However, most transcriptomic studies do not distinguish the relative contribution of transcription, RNA processing and RNA degradation processes to cellular homeostasis. Here we used metabolic labeling followed by massive parallel sequencing of newly transcribed and preexisting RNA fractions to simultaneously analyze RNA synthesis and decay in primary endothelial cells exposed to low oxygen tension. We found that the changes in transcription rates induced by hypoxia are the major determinant of RNA levels. However, degradation rates also had a significant contribution, accounting for 24% of the observed variability in total mRNA. In addition, our results indicated that hypoxia led to a reduction of the overall mRNA stability from a median half-life in normoxia of 8.7 hours, to 5.7 hours in hypoxia. Analysis of RNA content per cell confirmed a decrease of both mRNA and total RNA in hypoxic samples and that this effect was mimicked by forced activation of the Hypoxia Inducible Factor pathway and prevented by its interference. In summary, our study provides a quantitative analysis of the contribution of RNA synthesis and stability to the transcriptional response to hypoxia and uncovers an unexpected effect on the latter.

## INTRODUCTION

Eukaryotic cells show a remarkable regulatory flexibility that allows them to adjust to a constantly varying environment. Modulation of gene expression, consequence of changes at the transcriptional and post-transcriptional levels, plays a central role in the adaptation to environmental changes and stresses. However, quantitative information about the relative contribution of transcription to the changes in gene expression in response to specific signals is still incomplete.

The response to hypoxia, defined as the situation where oxygen consumption exceeds its supply, is a paradigmatic example of how gene expression shapes cell physiology to cope with environmental fluctuations. The Hypoxia Inducible Factors (HIFs) are transcription factors that play a fundamental role in this response. HIF is a heterodimeric transcription factor conserved in all metazoans (Iyer et al. 1998) that is composed of an oxygen-regulated alpha subunit (HIFα) and a constitutive expressed beta subunit (HIFβ) also known as Aryl receptor nuclear translocator (ARNT). There are three genes encoding HIFα subunits: HIF1A, EPAS1 (also known as HIF2A) and HIF3A. All these proteins belong to the PER-ARNT-SIM (PAS) subfamily of the basic helix-loop-helix (bHLH) family of transcription factors and they have a similar domain organization. HIFα and HIFβ subunits share an N-terminus bHLH domain that mediates the DNA binding and a central region or PAS domain that facilitates the heterodimerization (Jiang et al. 1997). Only the α subunits contain an oxygen dependent degradation domain (ODD) with two proline residues (Pro 402/ Pro 564 in HIF1A, Pro 405/ Pro 531 in EPAS1, and pro 492 in HIF3A) that are essential for the oxygen-induced HIF stabilization. Furthermore, HIF1A and EPAS1 present two transactivation domains NTAD and CTAD that can bind transcriptional co-regulators such as CBP/p300 and the cyclin dependent kinase 8 (CDK8) (Galbraith et al. 2013; Arany et al. 1996; Kasper et al. 2005; Dengler et al. 2014; Choudhry et al. 2014). The regulation of HIFα stability is mediated by a family of enzymes called EGL-nine homologues (EGLNs) that, in normoxic conditions, use molecular oxygen to hydroxylate specific proline residues in HIFα. The hydroxylation in one of these residues is sufficient for the recognition of HIF by the Von Hippel Lindau tumour suppressor protein (pVHL) that is the substrate recognition subunit of an E3-ubiquitin ligase complex. Thus, pVHL binding leads to HIFα ubiquitination, targeting it for degradation via proteasome. (Ivan et al. 2001; Jaakkola et al. 2001; Maxwell et al. 1999). In hypoxic conditions, HIF is stabilized and binds the RCGTG motif present in the regulatory regions of its targets genes and interacts with the transcriptional machinery to regulate their transcription.

The global gene expression pattern induced by hypoxia has been widely studied. (Xia and Kung 2009; Elvidge et al. 2006; Hu et al. 2003; Mimura et al. 2012; Salama et al. 2015; Scheurer et al. 2004; Warnecke et al. 2008; Weigand et al. 2012). However, these studies relied on RNA-seq and DNA microarrays that detect mRNA steady state-levels and therefore cannot distinguish between regulation of transcription and other mechanisms of post-transcriptional regulation. In this regard, apart from transcription, other processes such as RNA maturation and decay are under an exhaustive control and contribute to changes in gene expression. The relative importance of each process to determine the steady state expression levels of proteins has been studied in a few cellular systems, including the determination of basal levels of expression in fibroblasts (Schwanhausser et al. 2011) and the response of dendritic cells to lipopolysaccharide (Rabani et al. 2011). The conclusion of these and related studies is that transcription rate explains most of the observed variability in gene expression (Li et al. 2014; Rabani et al. 2011; Schwanhausser et al. 2011, 2013). However the generality of this conclusion, the extent to which the transcriptional changes mediated by HIF affect mRNA expression changes observed under hypoxia is unknown. In this regard, it has been demonstrated that, in addition to transcription, regulation of half-life of certain mRNAs has fundamental roles in hypoxia. For instance, the mRNA level of Vascular Endothelial Growth Factor A (*VEGFA*), a potent angiogenic factor, is regulated by hypoxia through both HIF-mediated transcriptional upregulation, as well as HIF-independent post-transcriptional regulation. *VEGFA* mRNA is an unstable transcript with a short half-life (60–180 min) in normoxia. However, under hypoxia *VEGFA* mRNA is stabilized by the binding of different RNA-binding proteins, such as HuR or hRNP-L, to AU-rich elements (AREs) present in its 3’-UTR. (Goldberg-Cohen et al. 2002; Levy et al. 1998, 1996; Shih and Claffey 1999). Other examples of transcripts whose half-lives are regulated by hypoxia through AREs include Erythropoietin (*EPO*) (Goldberg et al. 1991; Ho et al. 1995; Rondon et al. 1991) and tyrosine hydroxylase (*TH*) mRNAs. (Czyzyk-Krzeska et al. 1994). In addition, hypoxia has an important role in the regulation of miRNAs (Camps et al. 2014; Nallamshetty et al. 2013; Voellenkle et al. 2012), which in turn regulate mRNA stability and translation. Altogether, these results suggest a pervasive effect of hypoxia on mRNA stability, therefore, a global study of how hypoxia modulate mRNA half-life is needed to fully understand how cells adapt to low oxygen situations.

Herein, we determined the relative contribution of RNA transcription and stability to gene expression changes induced by hypoxia in primary human endothelial cells (Human Umbilical Vein Endothelial Cells, HUVEC) that are central actors in the induction of angiogenesis, a key adaptation response to hypoxia. Our results show that *de-novo* transcription explains 72% of gene expression changes observed upon low oxygen conditions, while half-life regulation underlies most of the remaining variability (24%). In addition, we found that the EGLN/HIF axis increases global RNA decay rates, leading to a 30% reduction of cellular mRNA as well as rRNA content. Thus, our work reveals an unexpected role of hypoxia on global RNA stability and provides quantitative data to understand the effect of hypoxia on gene expression.

## RESULTS

### Transcription rate is the major determinant of hypoxic mRNA levels

The transcriptional response to hypoxia has been studied in many different model systems at the genome scale (Xia and Kung 2009; Elvidge et al. 2006; Hu et al. 2003; Mimura et al. 2012; Salama et al. 2015; Scheurer et al. 2004; Warnecke et al. 2008; Weigand et al. 2012). However, in all these works mRNA levels were determined by means of RNA-sequencing or DNA microarrays and hence they could only assess the effect of hypoxia on steady state mRNA levels, which are determined by the combination of the relative rates of transcription and decay. Aiming to determine the contribution of these two processes in the response to hypoxia, we pulse-treated HUVEC for two hours with 4-thiouridine (4sU) to label nascent RNA and then characterized the pattern of newly transcribed mRNAs by affinity capture of the labelled transcripts followed by high-throughput sequencing (**Figure 1A**). Incorporation of 4sU into mRNA after a short pulse can be used as a proxy for transcription rate and comparison of changes in this nascent fraction to those in total RNA levels allows quantification of the role of mRNA stability on gene expression (Dolken et al. 2008; Schwanhausser et al. 2011). Hence, to determine the contribution of transcription to gene expression, we compared expression changes induced by hypoxia in the total mRNA fraction to those observed in the newly synthesized fractions (**Figure 1A**). As shown in **figure 1B**, taking a relatively relaxed threshold of FDR<0.05, only about one third of the genes found differentially expressed in the total RNA-fraction showed altered transcription (27% and 35% of the down- and upregulated genes respectively). However, this percentage gradually increased up to approximately 70% in both fractions when, in addition to the FDR, we used increasingly restrictive fold-change thresholds to consider genes as differentially-expressed in hypoxia (**Figure 1C**). This result suggests that transcription is mostly responsible for large expression changes in the total mRNA fraction, whereas changes of small magnitude are likely due to variations of mRNA half-lives. In addition, we observed a significant positive linear correlation between changes in gene expression observed in total and newly transcribed fractions (**Figure 1D**). By focusing on genes whose expression was significantly altered by hypoxia (FDR<0.01) in either total or newly transcribed mRNA fractions, we found that the transcriptional component (changes in newly synthesized mRNA) accounted for 72% of the observed variability in total mRNA (regression analysis, F1,2225=5756, p<10^-16^, R^2^2225=0.7212). In sum, these results suggest that transcriptional mechanisms largely determine gene expression changes in hypoxia.

**Figure 1.**
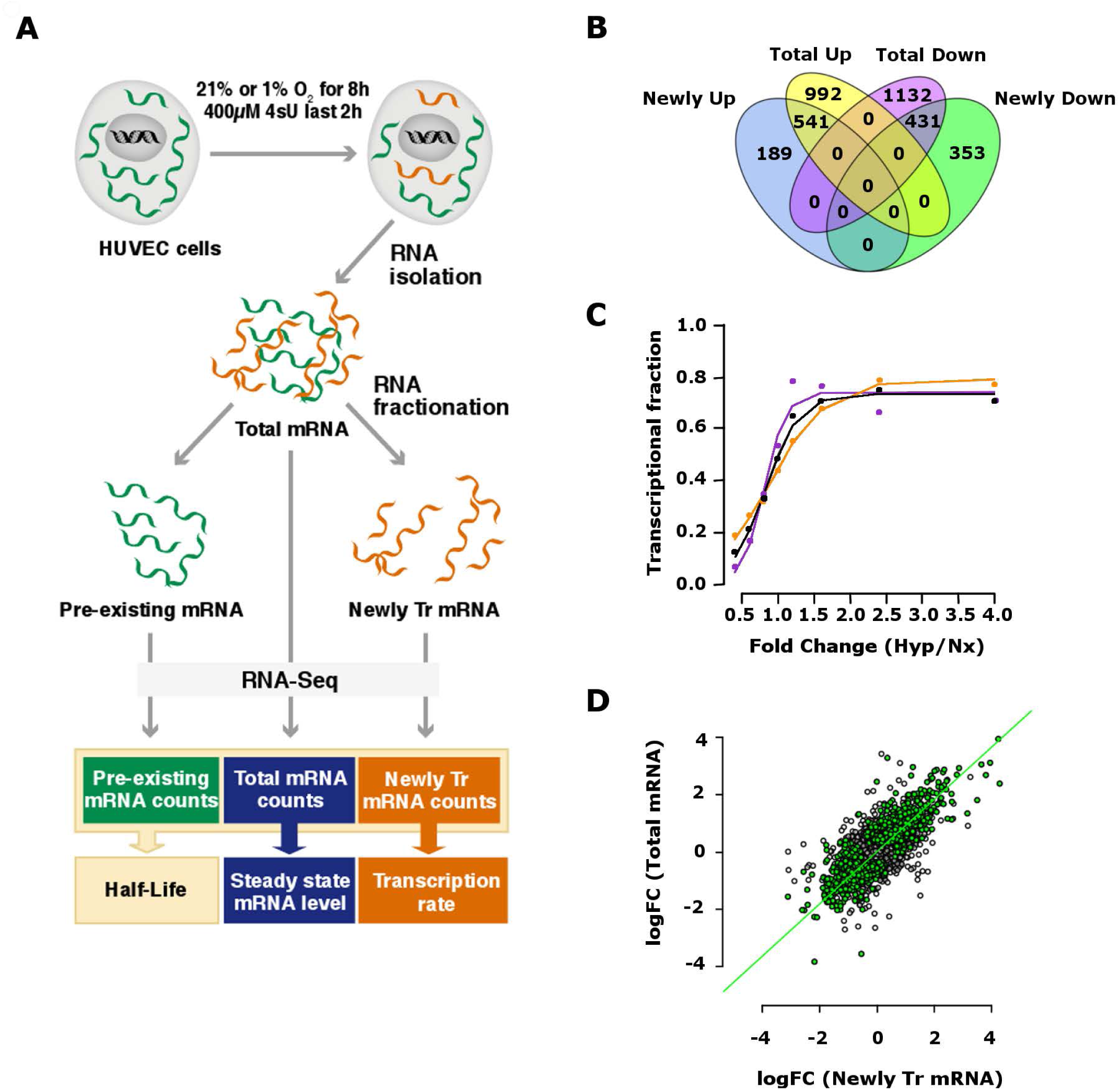
Hypoxia-induced changes in gene expression are predominantly controlled at the level of transcription. HUVEC were exposed to normoxia (21% oxygen) or hypoxia (1% oxygen) for 8 hours and pulse labelled with 4-thiouridine (4sU) during the last two hours of treatment. RNA was extracted from samples and an aliquot of it was fractionated into newly transcribed (4sU-labelled) and pre-existing (non-labelled) pools. For libraries preparation, poly-A RNA was purified from each of the three fractions (total, 4sU-labelled and pre-existing RNA), analysed by high-throughput sequencing and quantified by mapping reads to the exonic regions of genes. The results shown derive from three independent biological replicas, each using a different pool of HUVEC donors. **(A)** Schematic diagram depicting the experimental procedure and the information provided by each RNA fraction. **(B)** Venn diagrams showing the overlap between the genes found differentially overexpressed (FDR<0.05) in the total (Total Up) and newly transcribed (Newly Up) fractions and the differentially downregulated (FDR<0.05) in the total (Total Down) and newly transcribed (Newly Down) fractions. **(C)** The graph represents the proportion of genes in the total mRNA fraction that were also differentially expressed in the newly transcribed fraction for: all regulated genes (black line), significantly downregulated genes (purple line), and significantly upregulated genes (orange line). Genes were considered differentially expressed if, in addition of being significantly regulated (FDR<0.05), showed an absolute value of induction above the indicated threshold of logFC (x-axis). The magnitude of change induced by hypoxia was calculated for each gene as the absolute value of the logarithm of the ratio of the expression level in normoxia over hypoxia (logFC). **(D)** The magnitude of change (log2-fold change, logFC) induced by hypoxia in 4sU-labelled genes (“Newly tr mRNA”) was represented against the changes in the total mRNA fraction (“Total mRNA”). Genes significantly (FDR<0.01) regulated by hypoxia in either fraction are labeled in green. Regression analysis of significantly regulated genes showed that total mRNA level increased significantly with transcription (F1,2225=5756, p-value<<0.01).

### RNA stability is significantly affected by hypoxia

In spite of the major role of transcription, around 28% of the variability in total mRNA levels cannot be explained solely by this process. To further explore the contribution of non-transcriptional processes, we investigated the effect of hypoxia on mRNA half-lives by sequencing newly synthesized mRNA (4sU-labelled) in parallel with pre-existing (non-labelled) and total RNA factions from the same RNA sample (**Figure 1A**). This analysis allowed us to determine individual mRNA half-lives under normoxia and hypoxia (Dolken et al. 2008; Schwanhausser et al. 2011) (**Supplemental_table_S1**). We found that the distribution of mRNA half-lives in normoxic HUVEC was comparable to those reported for mouse fibroblasts (**Figure 2A**), with a median of 8.7h. In addition, half-life values determined by this technique were in good agreement with those obtained by measuring mRNA stability after inhibition of transcription by actinomycin D (**Figure 2B**; Pearson’s R48 =0.65, p<0.001, linear regression R^2^=0.77). Therefore, our determinations based on sequencing of the distinct mRNA fractions allowed for an accurate determination of mRNA half-lives. Next, we investigated whether hypoxia-induced changes on mRNA stability could account for the remaining variability in total mRNA levels that are not explained by transcription. A linear regression model including mRNA half-lives as an additional explanatory variable accounted for up to 96% of changes in total mRNA induced by hypoxia (multiple linear regression analysis, F2,1851=26110, p<0.0001, adjusted R^2^1851=0.9658). Thus, changes in mRNA half-life explain an additional 24% of the observed effects on gene expression and both variables, transcription and stability, jointly explain most of the variability in mRNA levels (**Supplemental_movie_s1**).

**Figure 2.**
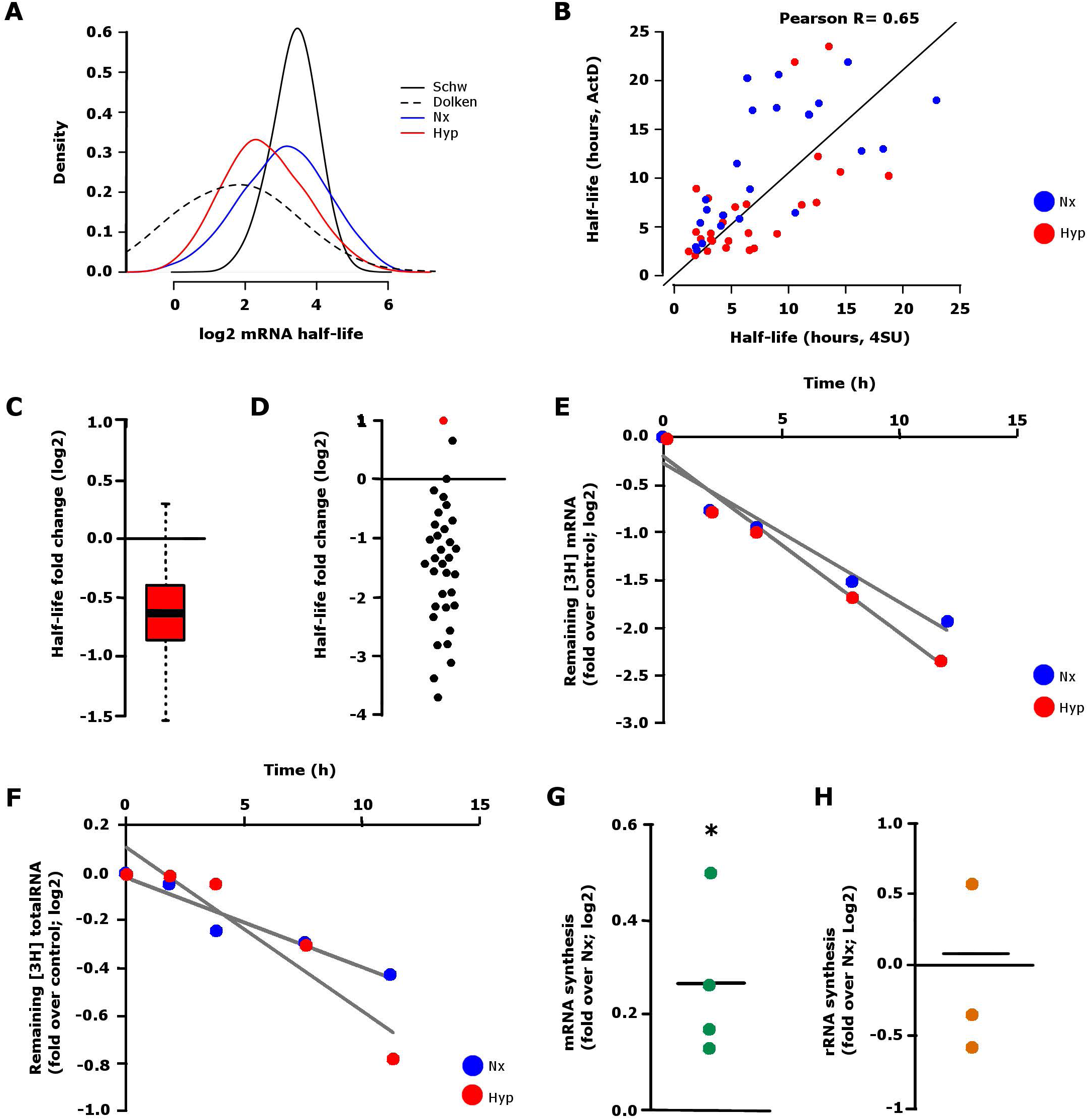
Decrease of mRNA and total RNA half-lives under hypoxic conditions. **(A)** Distribution of mRNA half-lives, determined as indicated in methods, in normoxic (blue) and hypoxic (red) HUVEC. The values published in (Dolken et al. 2008) (Dolken, dotted line) and (Schwanhausser et al. 2011) (Schw, black line) are overlapped for comparison. **(B)** Comparison of individual half-life values obtained by the high-resolution gene expression profiling experiment and the data derived from actinomycin D experiments. (F1,49=168, p<0.001,linear regression R^2^=0.77) **(C)** Distribution of changes in decay rates induced by hypoxia, as determined in the high-resolution profiling experiments, represented as the logarithm of the ratio of the half-life in hypoxia over normoxia. **(D)** HUVEC cells were grown overnight in normoxia or hypoxia and then treated with actinomycin D for 0h, 1h, 2h, 4h, 8, or 12 h. The graph represents the half-life of individual genes in normoxia versus hypoxia as log2 of the ratio. Each dot represents the median of three independent experiments. The red dot represents VEGFA. **(E and F)** HUVEC cells were pulsed with tritiated uridine for 2h and grown 16h in normoxia or hypoxia. Next, cells were treated with actinomycin D for 0h, 1h, 2h, 4h, 8h, and 12h and mRNA was purified from samples. Radioactive signal was used as readout of mRNA content to calculate half-life in normoxia (Nx, blue) and hypoxia (Hyp, red). The graph represents the logarithm of the ratio of the remaining radioactive signal in each time point over the radioactivity present at t=0 in polyA+ purified fraction **(E)** and in total RNA fraction **(F)**. **(G)** The amount of mRNA in the newly synthesized mRNA fraction (4sU-labelled) from the experiment described in Figure 1A was determined by spectrophotometry and represented as the log2 of the ratio of the values obtained in hypoxia over normoxia. Each point represents data from an independent experiment. The comparison of the observed ratios was significantly higher than expected from a sample with a mean log2-ratio of 0 (single sample Student’s t-test, p-value=0.049). **(H)** 47S rRNA precursor levels were determined by RT-qPCR in cells exposed overnight to normoxia or hypoxia and values normalized to 28S rRNA content. Each dot represents the ratio hypoxia over normoxia in an independent experiment. The observed ratios were not significantly different to the expected value of 1 (single sample Student’s t-test, p=0.8185).

In spite of its relatively minor role in the gene expression changes induced by hypoxia, the analysis of the effects of hypoxia on mRNA half-life revealed a modest, but significant (p<0.01 paired t-test), reduction of the overall mRNA stability from a median half-life in normoxia of 8.7 hours to a median half-life in hypoxia of 5.7 hours (**Figure 2A**; and **2C**). We confirmed this result by estimation of individual mRNA levels over time upon inhibition of transcription with actinomycin D under normoxia or hypoxia (**Figure 2D**). For these analyses we selected mRNAs showing different dependence on transcription in their response to hypoxia according to the 4sU analysis (**Figure 1B** and **Supplemental_table_S1**). Although the vast majority of mRNAs showed reduced half-life under hypoxia, our analysis also identified a few transcripts that were stabilized. Notably, among the latter was *VEGFA*, a transcript known to be stabilized in hypoxia (Goldberg-Cohen et al. 2002; Levy et al. 1998, 1996; Levy 1998; Shih and Claffey 1999). The pervasive effect on mRNA decay rates, suggested a general mechanism of RNA degradation rather than a sequence specific mechanism such as those mediated by miRNAs. Therefore, we explored if the effect of hypoxia on RNA decay rates were observed in unfractionated RNA; most of which consists in rRNA. With this objective, we pulse-labelled HUVEC cells with tritiated uridine and determined the decay of radioactive label in mRNA (**Figure 2E**) and total RNA (**Figure 2F**) over time after treatment with actinomycin D. As shown in **Figures 2E** and **2F**, the overall stability of total as well as mRNA was higher in normoxia than in hypoxia. These results pointed to a general increase in RNA decay rates rather than a specific effect on protein coding genes.

Although, reduced transcription rates could also potentially contribute to decreased RNA levels in hypoxic cells, we did not observe a decrease in 4sU incorporation in HUVEC cells exposed to 1 % oxygen compared to normoxic controls (**Figure 2G**). In fact, mRNA synthesis was slightly increased in hypoxia, suggesting very similar transcription rates in both situations.

In agreement, determination of the quantity of the 47S precursor ribosomal RNA (pre-RNA), which correlates with the rate of rRNA transcription (Cui and Tseng 2004), did not show significant differences between normoxia and hypoxia (**Figure 2H**).

In summary, these results suggest that hypoxia results in modest increase of RNA degradation without a significant impact in the RNA synthesis rates.

### Hypoxia-induced mRNA decay is HIF-dependent

The HIF axis is the best characterized signaling pathway in the response to hypoxia. Thus, as a first attempt to explore the molecular mechanism by which RNA degradation was increased in hypoxia, we investigated the effect of EGLN inhibitor Dimethyloxaloylglycine (DMOG) on total RNA mass. The EGLN family of enzymes act as the cellular oxygen sensors and function as the most apical elements of the HIF axis suppressing its activity in the presence of oxygen. As it is shown in **Figure 3A**, total RNA content was reduced in hypoxic conditions as well as in DMOG-treated cells compared with normoxia. Since HIF is the major target of EGLNs and a key player in the response to hypoxia, we next examined its involvement in this process by silencing the constitutive ARNT subunit. *ARNT* knock-down resulted in 36%-15% of control *ARNT* expression values and blocked the induction of the HIF target *EGLN3* (**Figures 3B** and **3C**). Importantly, *ARNT* knock-down significantly attenuated the RNA decay promoted by hypoxia (**Figure 3D**), suggesting that HIF activity is necessary in this process. To analyse if HIF stabilization was sufficient to promote RNA degradation, we infected HUVEC with lentiviral vectors that overexpressed constitutively active forms of *EPAS1* (EPASPP) or *HIF1A* (EPASPP, HIFPP) that had their oxygen-sensitive prolines residues substituted by alanine to prevent hydroxylation, rendering these isoforms constitutively active even in normoxia (Kondo et al. 2002). Additionally, we included mutant versions of these constructs where the bHLH motif was altered to prevent their capacity to bind DNA (EPASPPBH, HIFPPBH) (Kondo et al. 2002). As expected, the expression of EPASPP in normoxia resulted in a nine-fold increase of the *ADM* mRNA levels, whereas the expression of EPASPPBH had no effect (**Figures 3E** and **3F**). Similarly, the HIF1A-target *BNIP3* was strongly induced by overexpression of HIF1PP with intact bHLH domain (**Figures 3G** and **3H**). As it is shown in **Figures 3I** and **3J**, transduction of the constitutively active forms of both *EPAS1* and *HIF1A* promoted RNA decay in normoxia. In contrast, the over-expression of the mutants lacking a functional DNA binding motif did not reduce RNA content. Therefore, altogether these results suggest that HIF transcriptional activity is necessary and sufficient to induce RNA decay.

**Figure 3.**
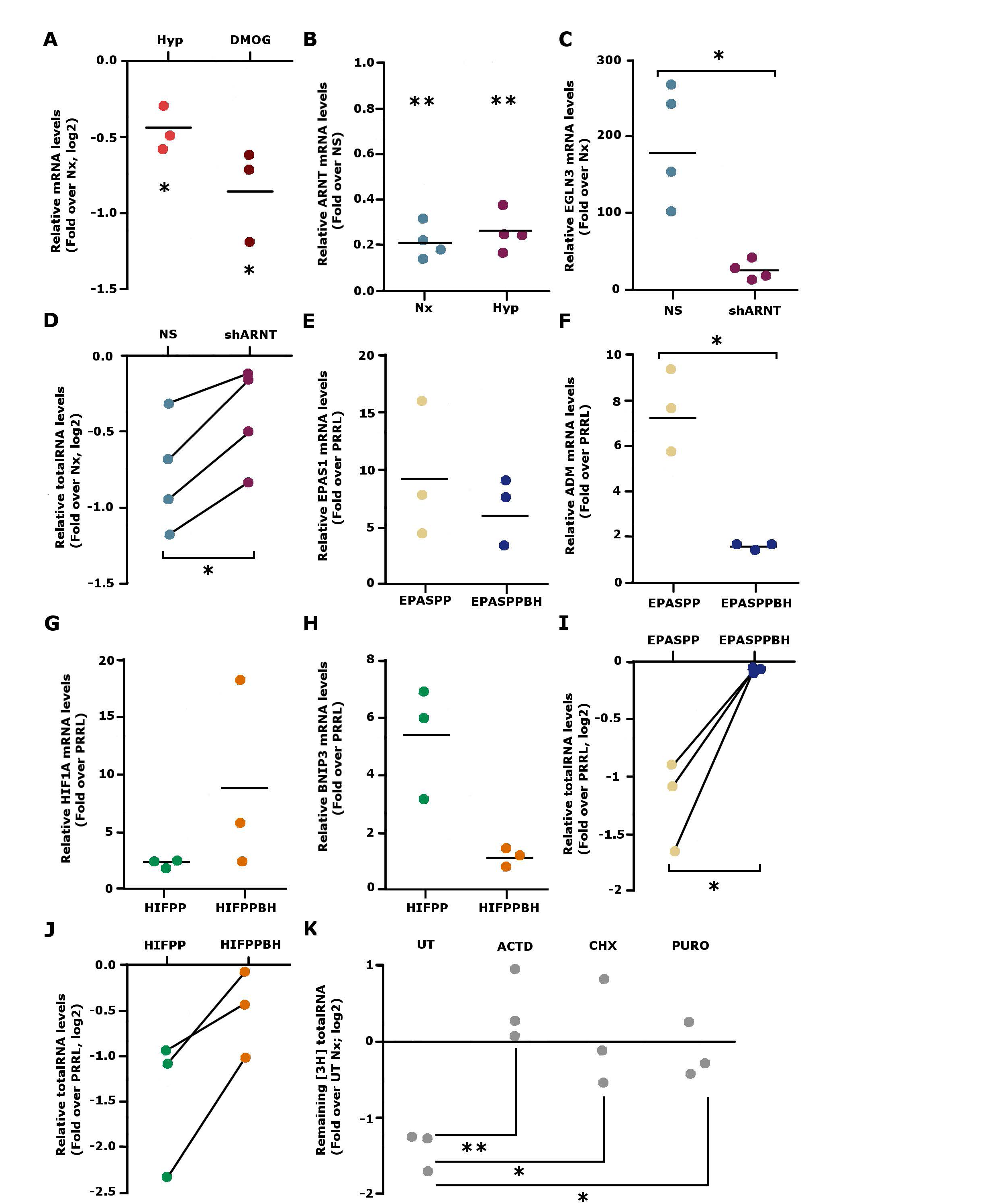
The increased mRNA decay in hypoxia depend on HIF transcriptional activity. **(A)** Total RNA and DNA were extracted from cells grown for 16 h in normoxia, hypoxia (Hyp) or DMOG (DMOG). PolyA+ RNA was purified from total RNA and then was normalized to DNA to calculate mRNA content per cell. The graph represents the ratio of RNA content in hypoxia or DMOG over normoxia. Each dot represents the ratio of RNA levels in hypoxia or DMOG over normoxia in an independent experiment. The observed ratios were significantly lower than expected from a sample with a mean ratio of 1 (single sample Student’s t-test, p-value=0.033 and p-value=0.041, respectively) **(B)** ARNT mRNA levels in HUVEC cells transduced with lentiviral particles expressing an shRNA to silence ARNT expression versus control (NS). The ARNT expression was on average 21% and 25% of that in control cells in normoxia and hypoxia respectively and significantly different to the expected value of 1 (single sample Student’s t-test, p=0.006 and p=0.007 for cells under normoxia and hypoxia respectively). **(C)** EGLN3 mRNA levels in HUVEC cells transduced with lentiviral particles expressing an shRNA to silence ARNT expression versus control (NS). The effect of hypoxia on EGLN3 induction was significantly attenuated in ARNT knockdown cells (p-value=0.017, paired Student’s t-test). **(D)** Total RNA and DNA were extracted from control cells (NS) and cells whose ARNT expression was silenced (shARNT), grown for 16 h in normoxia or hypoxia. Total RNA was normalized to DNA to calculate RNA content per cell. Each dot represents a determination from a single experiment and matched observations (derived from the same experiment) are represented joined by a line. The effect of hypoxia on RNA level was significantly attenuated in ARNT knockdown cells (p-value=0.014, paired Student’s t-test). **(E)** EPAS1 mRNA levels of HUVEC cells transduced with a constitutively stable form of EPAS1 (EPASPP) or a stable but transcriptionally inactive form (EPASPPBH) compared with the control (PRRL). The expression of mutant forms of EPAS1 was similar (p=0.43, paired Student’s t-test). **(F)** ADM mRNA levels of HUVEC cells transduced with a constitutively stable form of EPAS1 (EPASPP) or a stable but transcriptionally inactive form (EPASPPBH) compared with the control (PRRL). The effect of EPASPP on ADM expression was significantly higher than that of EPASPPBH (p=0.023, paired Student’s t-test). **(G)** HIF1A mRNA levels of HUVEC cells transduced with a constitutively stable form of HIF1A (HIFPP) or a stable but transcriptionally inactive form (HIFPPBH) compared with the control (PRRL). The expression of HIFPPBH was higher than that of HIFPP but the difference did not reached statistical significance (p=0.34, paired Student’s t-test). **(H)** BNIP3 mRNA levels of HUVEC cells transduced with a constitutively stable form of HIF1A (HIFPP) or a stable but transcriptionally inactive form (HIFPPBH) compared with the control (PRRL). The effect of HIFPP on BNIP3 expression was higher than that of HIFPPBH, but the difference did not reached statistical significance (p=0.079, paired Student’s t-test). **(I and J)** Total RNA and DNA were extracted from control cells and cells overexpressing a constitutively active EPAS1 (EPASPP) or HIF1A (HIFPP) and a constitutively active EPAS1 or HIF1A whose bHLH motive has been mutated (EPASPPBH, HIFPPBH). Total RNA was normalized to DNA to calculate RNA content per cell. The graph represents the ratio of RNA content per cell in overexpressing cells over control cells infected with a control lentivirus. Each dot represents a determination from a single experiment and matched observations (derived from the same experiment) are represented joined by a line. The expression of EPASPP resulted in RNA level significantly reduced as compared to control cells, (p-value=0.025, single sample Student’s t-test) and effect of each EPAS1 mutant on RNA level was significantly different (p-value=0.025, paired Student’s t-test). The expression of HIFPP resulted in RNA level reduced as compared to control cells, but did not reach statistical significance (p-value=0.08, single sample Student’s t-test) and effect of each HIF mutant on RNA level was not significantly different (p-value=0.23, paired Student’s t-test). **(K)** HUVEC cells were pulsed during 2 h with tritiated uridine and then treated with actinomycin (ACTD), cycloheximide (CHX) or puromycin (PURO) and exposed to hypoxia for 8h. The RNA was purified from samples in each condition and the remaining radioactivity was determined by scintillation counting. The graph represents the ratio of remaining radioactivity present in hypoxic samples over normoxia in untreated cells (UT) and cells treated with actinomycin D (ACTD), cycloheximide (CHX) or puromycin (PURO). Each dot represents a determination from a single experiment. A single factor ANOVA showed a significant difference among the four treatment groups UT, ACTD, CHX and PURO (F3,8= 8.487, p-value=0.007) and *a posteriori* Tukey test showed that all drug treatment means were significantly different than the control mean (p-value= 0.006, p-value=0.022 and p-value=0.047 for ACTD, CHX and PURO respectively).

The dependency on an intact bHLH domain suggested that the increased decay rate might be mediated by a HIF-target(s). To test this possibility, HUVEC cells were pulsed during 2 h with tritiated uridine, treated with actinomycin (ACTD), cycloheximide (CHX) or puromycin (PURO) and then exposed to hypoxia or normoxia for 8 additional hours. After these treatments, RNA was purified from samples and the remaining radioactivity was quantified by scintillation counting. As it is shown in **Figure 3K**, inhibition of transcription with actinomycin D or translation with cycloheximide or puromycin completely blocked RNA degradation in hypoxia. All these results suggest that a HIF target, whose transcription is induced by hypoxia, is responsible for the increased RNA decay rates.

## DISCUSSION

In the present work, we aimed to achieve a comprehensive and quantitative understanding of the gene expression changes that occur under adaptation of primary endothelial cells to hypoxic conditions. Previous studies focused on the analysis of the steady state mRNA levels, which are dictated by transcription and decay rates. Thus, we used a metabolic-labelling strategy to study the relative contribution of these processes on mRNA levels in response to hypoxia. In agreement with previous results on general mRNA regulation (Dolken et al. 2008; Schwanhausser et al. 2011; Rabani et al. 2011), we found that, although the relative contribution of transcription and decay rates is different for each individual gene, on average, transcription plays a dominant role on the changes in total mRNA induced by hypoxia (**Figure 1B**). Specifically, hypoxia-induced changes in transcription rates explain up to 72% of the variability in total RNA levels. Moreover, the fraction of differentially expressed genes shared by the total and newly transcribed fractions gradually decreases with more lenient thresholds of fold-change (**Figure 1D**). This result suggests that transcription is mostly responsible for large expression changes in the total mRNA fraction, whereas changes of small magnitude are mainly due to variations of mRNA half-lives. Given the role of miRNAs in fine-tuning expression levels at the post-transcriptional level and the impact of hypoxia on the expression of non-coding RNAs (Choudhry et al. 2014), it is tempting to speculate that the effects of hypoxia on mRNA half-lives could be mediated by miRNAs. Alternatively, this observation could just reflect that genes showing small changes in expression are harder to be detected as significant in the newly transcribed fraction. Finally, we also found genes whose changes in transcription did not correlate with changes in the total RNA fraction. One reason for this discrepancy is that the relative long half-life of mRNAs obscure the detection of changes in transcription by the determination of total expression levels.

Despite the major role of transcription, changes in half-life also contribute to gene regulation by hypoxia. According to our linear model, half-life explains nearly 24% of the variability on gene expression levels. A potential caveat of our analyses is that our determinations of transcription and decay rates, derive from the same 4SU-datasets and thus both variables are not independent, as formally required in multivariate linear regression. Nonetheless, the concordance of half-live values derived from the 4SU-labeling experiments and those derived from an independent technique based on inhibition of transcription with actinomycin D support the validity of our half-life estimations. As a generality, we observed a trend in the same direction for stability and changes in transcription. However, decay rates could have a high impact for the biology of certain genes. This is exemplified by *VEGFA*, a prototypical hypoxia-responsive gene, whose induction involves upregulation of transcription and decrease of decay rate (Levy et al. 1996; Shima et al. 1995). Changes in *VEGFA* mRNA stability allow for a fast response to hypoxia; while transcriptional regulation contributes as a later mechanism. Also, the half-life of *SLC23A2* (that encodes for the major transporter of ascorbate in endothelial cells) is dramatically reduced under hypoxia from 7.6 to 3 hours (**Supplemental_table_S1**). Ascorbate has been described to prevent HIF1A stabilization, (Kuiper et al. 2014, 2010) thus the rapid mRNA destabilization of *SLC23A2* could facilitate HIF induction under hypoxia. We did not investigate the mechanisms underlying these changes in mRNA stability, but they could be mediated by miRNAs or RNA binding proteins such as AUBPs, as it has been previously described for *EPO*; (Goldberg et al. 1991; Ho et al. 1995; Rondon et al. 1991) *VEGFA;* (Goldberg-Cohen et al. 2002; Levy et al. 1998, 1996; Levy 1998; Shih and Claffey 1999) or *EFNA3* (Pulkkinen et al. 2008).

In spite of the relatively minor role of mRNA decay on regulation of gene repression, we found that hypoxia resulted in modest (median reduction of 34-36%) but statistically significant reduction of the overall mRNA stability (**Figure 2A** and **2C**). This effect was confirmed by the determination of mRNA decay rates upon transcription inhibition with actinomycin-D (**Figure 2D**). We also found that, not only mRNA, but total RNA content per cell was also decreased in hypoxia (**Figure 2E** and **2F**). This result has two important implications. First, although hypoxia induces the transcription of a large number of miRNAs (Camps et al. 2014; Kulshreshtha et al. 2007) the global effect on both, mRNA and rRNA, makes miRNAs unlikely mediators of this process, and suggests a general mechanism of RNA degradation rather than a sequence specific mechanism such as those mediated by miRNAs. In search for a potential mechanism, we found that this effect is EGLN-HIF1A/EPAS1-dependent and relies on de-novo transcription and translation, pointing to a common target of HIF1A and EPAS1 as an effector for inducing RNA decay (**Figure 3**). In eukaryotes, the majority of mRNAs undergo decay through a pathway that is initiated by the shortening of the poly-A tail by deadenylases (PAN2–PAN3, CCR4–NOT and PARN). Therefore, by targeting some of the enzymes that are involved in the RNA-decay machinery, global RNA half-life could potentially be regulated. However, we did not observe hypoxic-induction of any of the genes encoding general RNA turnover proteins or endonucleases. In fact, we found decreased expression of several components of the exosome complex (*EXOSC2, EXOSC4, EXOSC6* and *EXOSC9*), required for general RNA degradation (data not shown). Nevertheless, we cannot rule out that hypoxia alters the protein level or activity of the enzymes involved in general RNA degradation.

Secondly, from the functional point of view, RNA stability is intimately coupled to translation. In fact, inhibition of translation in HUVEC cells resulted in a decrease in total RNA and mRNA levels (**Figure 4A**, **4B** and **4C**). Thus, the decreased RNA half-life observed in hypoxia could be a consequence of reduced translation observed under hypoxia, (Garneau et al. 2007). It has been long known that, in an attempt to save energy in oxygen-deprived cells, hypoxia diminishes protein synthesis by suppressing multiple key regulators of translation, including eIF2a, and the mammalian target of rapamycin (mTOR) (Liu et al. 2006; Wouters and Koritzinsky 2008). One of the mechanisms by which hypoxia inactivates mTOR and cap-dependent translation is by HIF-induction of *REDD1* in a TSC2-dependent manner (Wouters and Koritzinsky 2008). We have observed that translation is decreased in HUVEC cells upon oxygen deprivation or DMOG treatment (**Figure 4D**) and it correlates with dephosphorylation of the mTOR target rpS6 (**Figure 4E** and **4F**) that is partially prevented by EPAS1 silencing (**Figure 4G**). These results point out to a role of the EGLNs-HIF pathway in repressing translation in hypoxia in HUVEC. Interestingly, these are the same players that induce RNA decay upon oxygen deprivation (**Figure 3**). Although we cannot conclude that the increased RNA decay in hypoxia is a consequence of the inhibition of translation, our results show a good correlation between both processes. An attractive hypothesis is that in hypoxic cells mRNAs that are not being translated, together with the ribosomes, which are no longer needed for the translation of the hypoxic cell, are targeted for degradation; resulting in decreased half-life rates of both mRNA and rRNA. Alternatively, global reduction of half-life could contribute to repression of protein synthesis in hypoxia rather than being a consequence of it. Further work will be required to investigate all these possibilities

**Figure 4.**
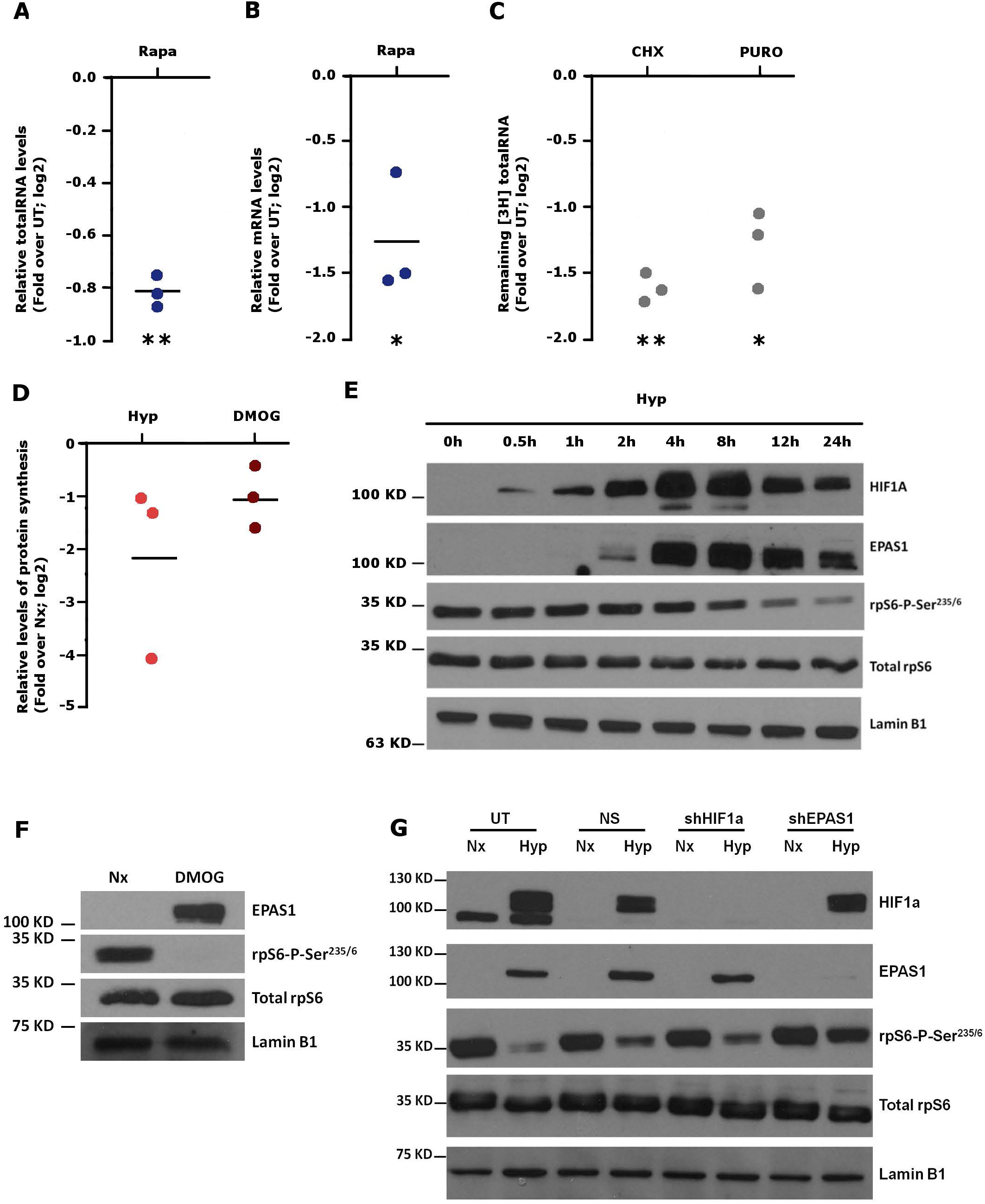
Effect of hypoxia and HIF on protein synthesis. **(A and B)** Cells were treated with rapamycin for 16h to inhibit translation. Then, RNA and DNA were extracted and polyA+ RNA was purified. Total RNA **(A)** and polyA+ RNA **(B)** were normalized to DNA content to calculate the amount of RNA per cell. The graph represents the ratio of RNA content per cell in rapamycin-treated cells over untreated HUVEC. Each dot represents a single experiment. The amount of total and messenger RNA per cell in rapamycin treated cells was significantly lower than in control cells (single sample Student’s t-test, p=0.002 and p=0.04 for total and messenger RNA respectively). **(C)** HUVEC cells were pulsed for 2h with tritiated uridine and then treated with cycloheximide, (CHX) or puromycin (PURO) to inhibit translation for 16h additional hours. Next, RNA was purified from samples in each condition and radioactivity was determined by scintillation counting. The graph represents the ratio of tritiated RNA present in treated over untreated samples. Each dot represents a single experiment. The amount of labelled RNA in CHX and PURO treated cells was significantly lower than in control cells. (single sample Student’s t-test, p=0.0015 and p=0.016 for CHX and PURO treated cells respectively) **(D)** HUVEC cells exposed to normoxia, hypoxia or normoxia in the presence of 500 µM DMOG for 16 hour and then pulse-labelled with S-35 methionine (200µCi/ml) for 30 minutes to determine translation rates. The graph represent the ratio of the radioactivity incorporated into proteins precipitated with TCA over total label in cell extracts. Each dot represents a single experiment. Both, hypoxia and DMOG, resulted in a strong inhibition of protein synthesis as compared to control conditions, but the differences did not reach statistical significance (single sample Student’s t-test, p=0.15 and p=0.1 for Hyp and DMOG treated cells respectively). **(E)** HUVEC were exposed to hypoxia for the indicated periods of time and the level of HIF1, EPAS1, phospho-S6 Ribosomal Protein (rpS6), total rpS6 and lamin B1 were determined by immunoblot. **(F)** HUVEC were treated with DMOG (DMOG) or vehicle (Nx) for 16 hours and the level of phospho-S6 Ribosomal Protein (rpS6), total rpS6 and lamin B1 were determined by immunoblot. **(G)** HUVEC cells transduced with lentiviral particles expressing a shRNA to silence HIF1A (shHIF1a) or EPAS1 (shEPAS1) expression, a control shRNA (NS) or mock infected (UT) were exposed to normoxia (Nx) or hypoxia (Hyp) for 16 hours. The level of HIF1, EPAS1, phospho-S6 Ribosomal Protein (rpS6), total rpS6 and lamin B1 were determined by immunoblot.

In addition to HUVEC, we have also observed decreased RNA content in hypoxia in other types of endothelial cells, such as Human Microvascular Endothelial Cells (HMVEC), but not in other cell types like Hela, HEK293T, human fibroblasts or mouse embryonic fibroblasts (data not shown). Thus, RNA degradation could be a specific response of endothelial cells to hypoxia. Alternatively, it could be a general response that is triggered at different hypoxic threshold depending on the cell type. In agreement, Hepa 1–6, (Johnson et al. 2008) C2C12 and NIH3T3 cells (Mekhail et al. 2006) but not HUVEC, blunt their RNA synthesis in response to very low oxygen concentrations or anoxia. Different cell types could diverge in the mechanisms employed to decrease RNA levels in hypoxia to inhibit translation.

In summary, our work supports that transcription is the major determinant of the relative abundance of transcripts in response to hypoxia. Notwithstanding, RNA stability significantly contributes to the changes in gene expression during adaptation to decreased oxygenation. In addition, we also uncovered an unexpected effect of hypoxia on total RNA content, due a HIF-mediated increase in global decay rates that could potentially contribute to fundamental global responses such as inhibition of translation. Finally, our data provides the first quantitative description of the effects of hypoxia on mRNA half-lives at global scale.

## MATERIALS AND METHODS

### Cell culture and treatments

Human Umbilical Vein Endothelial Cells (HUVEC) from pools of donors were purchased from Lonza (Lonza, C2519A) and grown in Endothelial Medium Bullet Kit (EGM-2, Lonza, CC-3162). For viral production, HEK 293T were maintained in Dulbecco’s modified Eagle’s medium (Gibco, 41966052) supplemented with 50 U/ml penicillin (Gibco, 15140122), 50 μg/ml streptomycin (Gibco, 15140122), 2 mM glutamine (Gibco, 25030123) and 10% (v/v) fetal bovine serum (Gibco, 10270106). All cells were grown at 37°C and 5% CO2 in a humidified incubator and tested regularly for mycoplasma contamination. For hypoxia treatment, cells were grown at 37°C in a 1% O2, 5% CO2, 94% N2 gas mixture in a Whitley Hypoxystation H35 (Don Withley Scientific). Alternatively, hypoxic conditions were mimicked using 500 µM Dimethyloxaloylglycine (DMOG, Sigma, D3695). Actinomycin, cicloheximide, puromycin and rapamycin purchased from sigma, were used at 5 µg/ml, 10ug/ml, 2ug/ml and 20nM respectively.

### Metabolic labelling with 4-thiouridine and purification of newly synthesized mRNA

We used the protocol described by Dolken et al, and Schawnhausser et al (Schwanhausser et al. 2011; Dolken et al. 2008) that is represented in the Figure 1A. Briefly, exponentially growing HUVEC cells were exposed to 21% (normoxia) or 1% (hypoxia) oxygen for 8 hours and pulse labelled with 4-thiouridine (400 µM, 4sU, Sigma, T4509) during the last two hours of treatment. After treatment, total cellular RNA was isolated from cells using TRI-reagent (Ambion, AM9738). 100 µg of total RNA was subjected to a biotinylation reaction to label the newly transcribed RNA containing the 4sU moiety (Pierce, EZ-Link Biotin-HPDP, 21341). Then, RNA was purified using Ultrapure TM Phenol:Chloroform:Isoamylalcohol (Invitrogen, 15593 031) and labelled RNA was isolated from the total RNA by affinity chromatography using streptavidin coated magnetic beads (μMacs Streptavidin Kit; Miltenyi, 130074 101).

### High-throughput sequencing and bioinformatics analysis

Libraries of total RNA, labelled RNA and unlabelled RNA were prepared using the standard protocol for messenger RNA sequencing (Illumina, TruSeq Stranded mRNA) and sequenced on HiSeq2000 instrument (Illumina) according to the manufacturer’s protocol. The resulting reads were aligned to the GRCh37/hg19 assembly of the human genome using TopHat (Trapnell et al. 2009) with the default parameters. Finally, the gene expression level was calculated as the number of reads per gene, computed using HTSeq (Anders et al. 2015) and gene features as defined in the GRCh37.75 release of the human genome (gtf file). Differential gene expression analysis was performed with the Bioconductor (Gentleman et al. 2004) edgeR package (Robinson et al. 2010) for the R statistical Software (http://www.R-project.org/). The raw reads and processed data derived from RNA seq experiments are available at NCBI’s Gene Expression Omnibus (Barrett et al. 2009) and are accessible through the GSE89831 GEO Series accession number. All computations were performed using R software package (http://www.R-project.org/).

### mRNA half-life determination

Individual mRNA half-lives were calculated as described in (Dolken et al. 2008). For each mRNA half-life was computed according to the following equation:

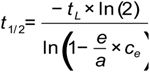

Where *e* and a represents the amount of newly transcribed RNA and total mRNA respectively, expressed as the normalized counts from the RNA-seq experiment. *tL* is the labelling time and *ce* is a correction factor estimated from slope of the linear regression fit of the u/a and e/a values for all the genes expressed in HUVEC according to the formula:

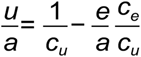

In this equation *u* represents the amount of pre-existing mRNA expressed as the normalized counts from the RNA-seq experiments. The *Ce* and *Cu* correction factors can be determined by the estimate of the slope and intercept of the linear fit of the expression level of all three fractions (total, newly transcribed and pre-existing). All computations were performed using R software package.

#### RNA extraction and qPCR

RNA was extracted and purified using the RNeasy Mini Kit (Qiagen; 74106) following manufacturer’s instructions. For quantitative-RT PCR analysis 1 µg of total RNA of each sample was reversed-transcribed to cDNA (Transcriptor First Strand cDNA Synthesis kit, Roche 04379012001), cDNA was diluted 1:20 and used as template for amplification reactions, carried out with the Power SYBR green PCR Master Mix (RNeasy Mini Kit Applied Biosystems; 4367659) following manufacturer’s instructions.

PCR amplifications were carried out in a StepOne Realtime PCR System (Applied Biosystems; 4376357). Data were analyzed with StepOne software and expression levels were calculated using ΔΔCT, using β-actin as reference. Primers sequences used for qPCR analysis are provided in **Table 1**.

**Table 1.**
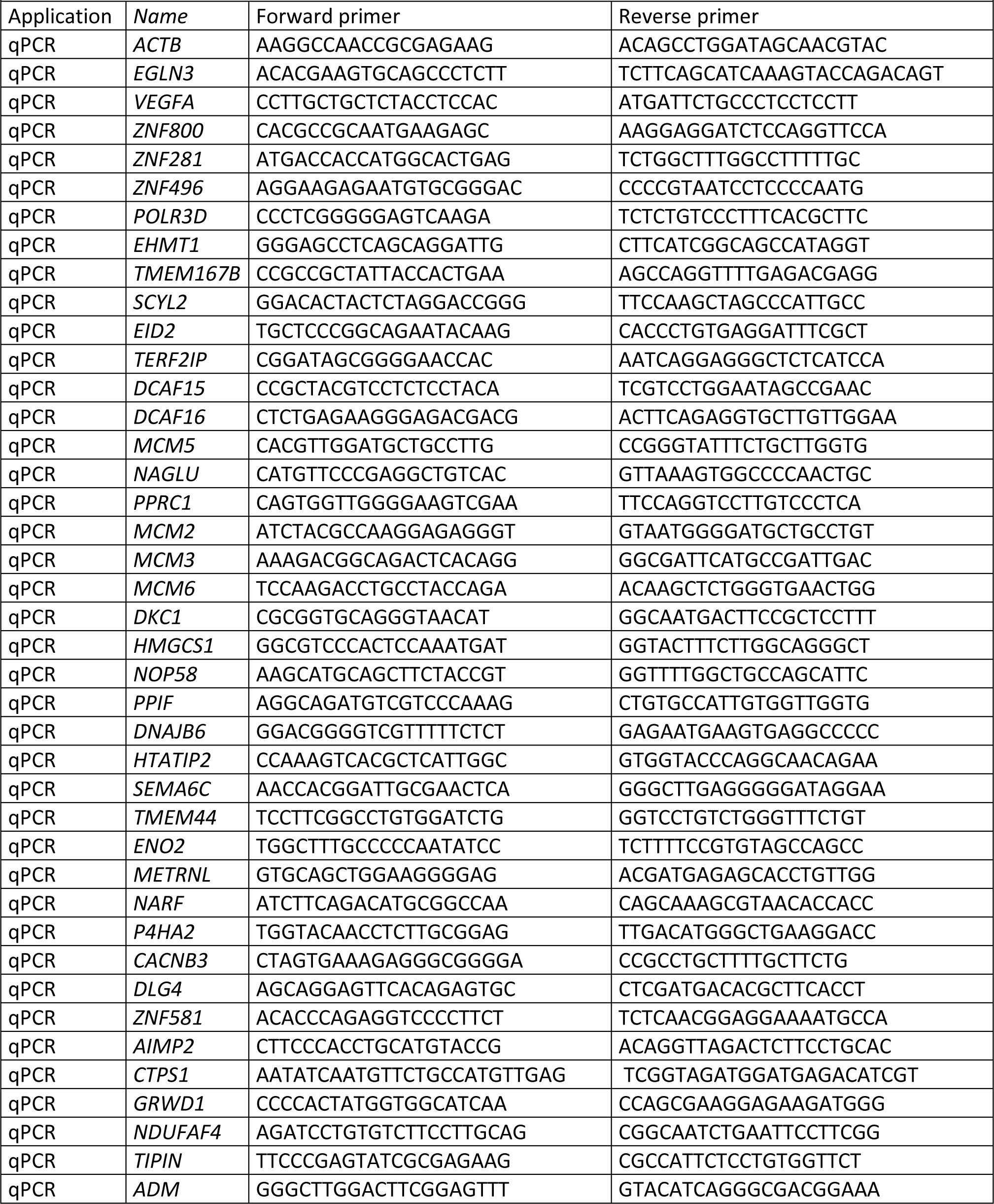
Primers used in this study

#### Individual mRNA half-life calculation using actinomycin D

HUVEC cells were seeded in 6 cm^2^ plates at a density of 4 x 10^5^ cells/well 8 hours prior to experiments. To determine the effect of hypoxia on mRNA decay rates, cells were exposed to normoxia or hypoxia for 16 hours. Then, treated with 5 µg/ml actinomycin D (Sigma; A9415), a general transcription inhibitor, and RNA was isolated 0h, 1h, 2h, 4h, 8h and 12h after transcription inhibition. Cells were kept at the same oxygen tension for the whole duration of the experiment.

The relative amount of specific mRNAs were determined by quantitative RT-PCR. Their decay constant (k_decay_) was obtained from the slope of the regression line obtained by the representation of the fraction of mRNA remaining at each time point:

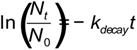

N_t_/N_0_ is the remaining fraction of mRNA at time point *t* after actinomycin D addition. Finally, the half-life for each mRNA was calculated from its decay constant as:

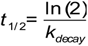

#### Pulse-labelling of HUVEC and HMVEC cells with tritiated uridine

HUVEC cells were seeded in 6 cm^2^ plates (4 x 10^5^ cells/well) and 8 hours after plating were pulsed for 2 hours with tritiated uridine (Uridine [5,6-^3^H]; Perkin Elmer; NET367001MC) at final concentration of 2 µCi/ml. After labeling, cells were washed twice with Phosphate Buffered-Saline (PBS), and incubated in non-radioactive complete culture medium under normoxia or hypoxia for 16 hours. The radioactivity present in Poly-A^+^ and total RNA fractions was measured using Liquid Scintillation (Optiphase Hisafe II; Perkin Elmer, 1200-436) and the Liquid Scintillation Analyzer (Perkin Elmer; TriCarb 2800TR). For RNA half-life determination, 5 µg/ml actinomycin D was added to block transcription and RNA was isolated at different time points after treatment (0h, 2h, 4h, 8h and 12 hours) followed by purification of Poly-A^+^ RNA using the Oligotex-DT mRNA Mini Kit (Qiagen; 50970022). Global mRNA half-life (*t* _1/2_) was calculated using the radioactive label to determine the ratio N_t_/N_0_.

#### rRNA synthesis determination

The determination of the 47S precursor ribosomal RNA (pre-RNA), was performed by qPCR and normalized by 28S using the primers described in **Table 1**

#### Calculation of total RNA and mRNA mass per cell

HUVEC cells were seeded in 6 cm^2^ plates at a density of 4.8 x 10^5^ cells per plate and were exposed to normoxia, hypoxia or normoxia in the presence of 500 µM DMOG for 16 hours. Then DNA and RNA were isolated from each sample and determined by spectrophotometry (NanoDrop ND-1000 instrument, Thermo Scientific). We calculated RNA mass per cell considering that human cells contain 6,6 pg of DNA per cell.

#### Lentiviral production

Lentiviral vector pGIPz (Open Biosystems) was used to silence the mRNA expression of *ARNT* (V3LHS_393533), *HIF1A* (V2LHS_132150), *EPAS1* (V2LHS_113753) and *TSC2* (V3LHS_642078), as well as a non-silencing lentiviral shRNA Control (NS; RHS4346). For the overexpression of HIF mutants stabilized in normoxia, we used PRRL lentiviral vectors containing *HIF1A* and *EPAS1* mutant sequences under cytomegalovirus (CMV) promoter (Kondo et al. 2002). These vectors contain the ODD prolines residues converted to alanine for maintaining HIF constitutively active in normoxic conditions (pRRL-HIF1A-P402A;P564A-IRES-eGFP and pRRL-EPAS1-P405A;P531A-IRES-eGFP). In addition, we used other HIF overexpressing vectors whose bHLH motif had been mutated to prevent DNA-binding (pRRL-HIF1A-P402A;P564A-bHLH* - IRES-eGFP and pRRL-EPAS1-P405A;P531A-bHLH*-IRES-eGFP). Lentivirus were produced and titrated in HEK293T cells as previously described (Punzon et al. 2004). For transduction of HUVEC cells, lentiviruses were used at a multiplicity of infection (MOI) of 1–2 for 8 h, resulting in more than 95% transduced (GFP-positive) cells 72 h after infection.

### Western blot

Cells were harvested in RIPA buffer (50mM Tris-HCl pH 8, 150 mM NaCl, 0,02% NaN3, 0,1% SDS, 1% NP-40, 1% Sodium Deoxycholate) containing protease inhibitors (Complete ULTRA tablet; Roche; 06538304001). Protein concentration was quantified using BioRad DC protein assay (BioRad, 5000112) and 40 μg of each sample was resolved in 10% SDS-polyacrilamide gels. Proteins were then transferred to Polyvinylidene difluoride (PVDF) membrane (Immobilon®-P, Millipore, IPVH100010). Membranes were blocked with 5% non-fat dry milk in TBS-T (50 mM Tris-HCl pH 7.6, 150 mM NaCl, 0.1% Tween-20) and incubated with the corresponding antibody (HIF1A: abcam; ab2185; EPAS1: Novus; NB100-122, LaminB1: Santa Cruz, sc-6216, Phospho-S6 Ribosomal Protein (Ser235/236), Cell Signaling, 4858; and S6 Ribosomal Protein, Cell signaling, 2217).

### Methionine S-35 pulse-labelling of HUVEC cells

HUVEC cells were seeded in 6 cm^2^ plates at a density of 4.8 x 10^5^ cells per plate and exposed to normoxia, hypoxia or normoxia in the presence of 500 µM DMOG for 16 hours. Then, cells were washed with DMEM lacking methionine and containing 10% of dialyzed FBS to eliminate the excess of methionine and 25mM Hepes. Medium was removed and 0.5ml of DMEM with ^35^S-35 methionine (200µCi/ml) was added, then cells were incubated at 37° 30 minutes. Cells were washed with cold PBS and recovered by cell-scrapping and centrifugation and precipitated with 10% TCA in glass fiber filter disks. To calculate the ratio of TCA-precipitated label relative onto total radioactivity, we measured the amount of radioactivity of each disk by scintillation counting and compared with the total radiolabeled cell suspension fraction obtained before precipitation.

## AKNOWLEDGMENTS

This work was supported by Ministerio de Ciencia e Innovación (Spanish Ministry of Science and Innovation, MICINN) [SAF2014-53819-R to Luis del Peso and Benilde Jimenez; SAF2017-88771-R to Luis del Peso and Benilde Jimenez].

