## Supplementary figures and images for "Metabolic labelling of RNA uncovers the contribution of transcription and decay rates on hypoxia-induced changes in RNA levels"

### Supplemental Movie 1

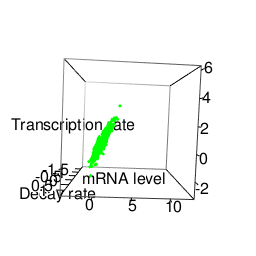
